## Appendix for "Uncovering the Organization of Neural Circuits with Generalized Phase Locking Analysis"

### Contact for reagent and resource sharing

### Experimental model and participant details

The neural data used in this study were recorded from the ventrolateral prefrontal cortex (vlPFC) of one anaesthetised adult, male rhesus monkey (macaca mulatta) by using Utah microelectrode arrays [Blackrock Microsystems [75]] (more details on these experiments are provided in a previous study exploiting this data by Safavi et al. [102]). All experiments were approved by the local authorities (Regierungspräsidium) and were in full compliance with the guidelines of the European Community (EUVD 86/609/EEC) for the care and use of laboratory animals.

### Method details

#### Simulation of phase-locked spike trains

We use simulated phase-locked spike trains and noisy oscillations as a toy model to demonstrate the potential applications of GPLA. The core principles of simulations used in both Figure 2, 4 and 3 are explained in the following paragraphs and the specializations used for individual figures are provided at the end (also summarized in table 2).

**Table 2**

| Figure num. | Osc. type | Num. of oscillatory components | Equations |
|---|---|---|---|
| Figure 2 | Transient | 1 | 16,34 |
| Figure 3 A-C | Transient | 1 | 16,34 |
| Figure 3 D-I | Sustained | 1 | 34,35 |
| Figure 4 A-G | Sustained | 5 | 17,34 |
| Figure 4 H | Sustained | 1-10 | 17, 34 |

For generating phase-locked spike trains, we adopt the method introduced in [4]. As the model has already been described elsewhere, we restrict ourselves to a brief explanation. We sample the spike times from an inhomogeneous Poisson process with rate $\lambda(t)$,

$$\lambda(t) = \lambda_0 \exp\left(\kappa \cos(2\pi f t - \varphi_0)\right) , \tag{34}$$

where $\varphi_0$ is the locking phase of the spikes with respect to the oscillation, $\kappa$ is the concentration parameter of the spikes around the locking phase, that specify the strength of coupling between spikes and the oscillation, $\lambda_0$ is propositional to the average firing rate over time ($\lambda_0 I_0(\kappa)$ is the average firing rate), and $f$ is the frequency of oscillatory modulation of the spike trains.

Furthermore, we can also derive an analytical expression for the complex-valued PLV to be used as ground truth PLV (used in Figure 3),

$$PLV^* = e^{\mathbf{i}\varphi_0} \frac{\int_0^\pi \cos(\theta)\exp(\kappa\cos(\theta))d\theta}{\int_0^\pi \exp(\kappa\cos(\theta))d\theta} = e^{\mathbf{i}\varphi_0}\frac{I_1(\kappa)}{I_0(\kappa)} , \tag{35}$$

where $PLV^*$ indicate the ground truth value, and the $I_k$'s denoting the modified Bessel functions of the first kind for $k$ integer (see e. g. Abramowitz et al. [2, p. 376]):

$$I_k(\kappa) = \frac{1}{\pi}\int_0^\pi \cos(k\theta)\exp(\kappa\cos(\theta))d\theta\,. \tag{36}$$

For the simulation used in Figure 4, we construct the LFP by superimposing $N_{osc} \leq 10$ oscillatory components $O_j(t) = e^{2\pi \boldsymbol{i} f_j t}$, $j \in \{1, \dots, N_{osc}\}$ that the frequency of oscillations are limited in range of $[f_{min},\, f_{max}]$. Each LFP signal is a weighted sum of these oscillatory components. We can represent these weights in a $(n_c \times N_{osc})$-variate matrix (we call it *mixing matrix*, and denote it by $W$), where each row of the mixing matrix indicate the weights for the corresponding LFP channel. Thus, the synthesized multichannel LFP ($\boldsymbol{\Psi}(t) = \{\psi_l(t)\}_{l=1,\dots,n_c}$) can be written as the product of the mixing matrix ($W$) and the oscillatory basis ($\boldsymbol{O}(t) = \{O_j(t)\}_{j=1,\dots,N_{osc}}$),

$$\boldsymbol{\Psi}(t) = W\boldsymbol{O}(t) + \boldsymbol{\eta}(t)\,, \tag{37}$$

where $\boldsymbol{\eta}(t)$ is additive white noise on both real and imaginary parts.

In this simulation, the frequency of the oscillatory components range from 11Hz to 15 Hz, and the mixing is the following,

$$W = \begin{bmatrix} \boldsymbol{w}_d & \boldsymbol{w}_0 & \cdots & \cdots & \boldsymbol{w}_0 \\ \boldsymbol{w}_0 & \boldsymbol{w}_d & \boldsymbol{w}_0 & \cdots & \boldsymbol{w}_0 \\ \boldsymbol{w}_0 & \boldsymbol{w}_0 & \ddots & \ddots & \boldsymbol{w}_0 \\ \boldsymbol{w}_0 & \ddots & \ddots & \ddots & \boldsymbol{w}_0 \\ \boldsymbol{w}_0 & \cdots & \cdots & \boldsymbol{w}_0 & \boldsymbol{w}_d \end{bmatrix}. \tag{38}$$

where $\boldsymbol{w}_d = w_d \mathbf{1}_{N_g}$ and $\boldsymbol{w}_0 = w_d \mathbf{0}_{N_g}$, $w_d = 1$ and $w_0 = 0.1$ ($\mathbf{1}_{N_g}$, is a $(N_g \times 1)$-variate all-one column vector). This simple structure of the mixing matrix (which is close to a block diagonal matrix) implies that each LFP channel contains one dominant oscillatory component with a specific frequency (as in each row, there is only one oscillatory component with large coefficient $w_d$ and a specific frequency).

For the simulation used in Figure 3A-C, oscillations originate from a single oscillatory source, but in order to make transient rather sustained oscillations, they were multiplied by a Gaussian window (with the size of 20 cycles of oscillation) around random events. The timing of transitory events was governed by a homogeneous Poisson process. Moreover, the spiking activities are phase-locked to the phase of the oscillations as the spike times were drawn from an inhomogeneous Poisson process with the rate specified in Equation 34. For the rest of the simulations in Figure 3 a single sustained oscillation has been used.

### Simulation of hippocampal sharp wave-ripples

The model was introduced and described in Ramirez-Villegas et al. [96]. We thus restrict ourselves to a brief explanation of the characteristics that are the most relevant to GPLA analysis.

#### Network architecture

We use a model of part of the hippocampal formation, accounting for the dynamics of CA1 and CA3 subfields during non-Rapid Eye Movement (non-REM) sleep. Cells of each subfield consist of 150 units (135 pyramidal neurons and 15 interneurons), arranged on a one dimensional array, along the x-axis. The connectivity of CA3 is characterized by strong recurrent excitatory auto-associational pyramidal-pyramidal connections, together with pyramidal-interneuron connections and short-range and interneuron-pyramidal synapses. In contrast, CA1 connectivity is implemented as a "feedback and reciprocal inhibition" circuit, including only pyramidal-interneuron, interneuron-pyramidal, and interneuron-interneuron synapses, all located in their peri-somatic region. Both excitatory and inhibitory populations in CA1 additionally receive inputs from afferent CA3 excitatory neurons (see Figure 6A for the schema of the model).

### Cell dynamics

Each neuron is modeled with two compartments: dendritic and axosomatic, the dynamics of each follows a Hodgkin-Huxley type (conductance-based) equation [90, 118]. Notably, they include a non-linear slow dendritic calcium channel responsible for the bursting activity.

### Computation of the laminar LFP profile

The procedure for computing laminar LFP profiles was also described in Ramirez-Villegas et al. [96]. Briefly, the trans-membrane current of each compartment of each cell is modeled as a line source [104] that were placed with the equal distance across the horizontal in a Stratum Pyramidale (SP) of 100$\mu$m thickness, with an axosomatic compartments height of 80$\mu$m for both pyramidal neurons and interneurons [118]. Total dendritic arbor height of pyramidal cells was 200$\mu$m, corresponding to the CA1 Stratum Radiatum (SR). LFPs were captured through two multi-channel electrodes (mimicking laminar probes), each with 16 recording sites disposed along the vertical axis (denoted by $z$), 20 $\mu$m apart covering the simulated axosomatic and apical dendritic fields of CA1 and CA3. Each electrode crosses the corresponding linear cell arrangement (perpendicularly) in its middle.

The extracellular medium is modeled as a uniform and isotropic ohmic conductor with resistivity $\rho = 333\Omega$cm. The potential in the extracellular medium is governed by the Poisson equation $\nabla^2 \phi = \frac{1}{\sigma}\frac{d\xi}{dt} = -\frac{I_t}{\sigma}$, where $\sigma = \frac{1}{\rho}$ is the conductivity of the extracellular space $\left[\frac{\text{S}}{\text{m}}\right]$. With these assumptions, the extracellular potential $\phi\left(y_0, r, t\right)$ at the algebraic depth $z_0$ and a radial distance $r$, measured over the compartment's length limits ($z_1$ and $z_2$, respectively indicate the algebraic depth and the bottom of the top of the cylindrical compartment with length $L = z_2 - z_1$) can be computed by

$$\phi\left(z_0, z_1, z_2, r, t\right) = \frac{1}{4\pi\sigma L}\int_{z_1}^{z_2} \frac{I\left(t\right)}{\sqrt{\left(z - z_0\right)^2 + r^2}} \mathrm{dz} = \frac{1}{4\pi\sigma}\frac{I\left(t\right)}{L} \ln\left[\frac{\sqrt{\left(z_1 - z_0\right)^2 + r^2} - \left(z_1 - z_0\right)}{\sqrt{\left(z_2 - z_0\right)^2 + r^2} - \left(z_2 - z_0\right)}\right]. \tag{39}$$

after solving the integral with standard procedures. Accounting for the contribution of all compartments and cells, the total extracellular potential $\phi_{\text{tot}}\left(z_0, t\right)$ at a given depth $z_0$ is

$$\phi_{\text{tot}}\left(z_0, t\right) = \sum_i \sum_j \phi_{i,j}\left(z_0, z_{1,i,j}, z_{2,i,j}, r_{i,j}, t\right)\,, \tag{40}$$

where $\phi_{i,j}$ is the potential generated by the total transmembrane current of the $j^{th}$ compartment of the $i^{th}$ cell, located at radial distance $r_{i,j}$ from the electrode.

Note that since the neuron models considered in this work are two-compartmental, charge conservation within the cell implies that the total absolute somatic transmembrane currents equal the absolute of the total dendritic transmembrane currents (which also follows the charge conservation principle), leading to a dipolar distribution of the LFP contribution for each cell.

Equations 39-40 describe the way LFPs are simulated (as the low-frequency parts of the extracellular potential) in the original biologically realistic model that generated the LFP data we use for GPLA analysis. We also exploit it to provide an approximate LFP laminar profile for the population of pyramidal cells based on this equation, by injecting the same constant current to all cells and compartment of the linear arrangement, but having opposite signs for the axosomatic and dendritic compartment to respect charge neutrality and thus the dipolar structure (Figure 6D (broken line)).

#### Neuron exclusion criterion for GPLA

To reduce the small sample bias caused by a low number of spike events, we only use neurons that had a minimum average firing rate of 3 Hz firing. Nevertheless, using all neurons did not change the results significantly. For instance, in Figure 6G (in contrast to Figure 6A-F and H where excluded neurons based on their firing), to be compatible with the neural mass model we did not exclude any neuron.

### Analytical neural field modeling of spike-field coupling

In order to justify and interpret our approach, we use a rate-based neural field model. Units are grouped in populations according to their cell-type on spatial localization. Spiking activity of a specific population $p$ at possibly multidimensional location $x$ is represented by its average spike rate $\lambda_p(x,t)$. Simultaneously, the LFP $L(X,t)$ is recorded at locations reflected by possibly different coordinates $X$.

#### Rate model of circuit dynamics

We follow classical neural field models, stating that the rate of each population evolves as a monotonous function of the membrane potential, itself controlled by post-synaptic currents (PSCs). Dynamics of the membrane potential $V_p$ of each population $p$, is assumed to be governed by the following differential equation:

$$\frac{dV_p}{dt}(x,t) + \tau_p V_p(x,t) = \alpha_p \eta(x,t) + \sum_k \nu_{p\leftarrow k} s_k(x,t)\,, \tag{41}$$

where $\eta$ represents the post-synaptic current generated by the external input (for which no spiking activity is available), $s_k$ the normalized[6] post-synaptic current from the afferent population $k$, whose effect on target population $p$ is scaled by synaptic strength $\nu_{p\leftarrow k}$.

The relationship between the normalized post-synaptic current at location $x$ and spiking activity of the afferent population activity is modeled by spatio-temporal integration [128, 109, 55], possibly taking into account the propagation speed $v_0$ along the axons

$$s_k(x) = \int c_k(x,X)\lambda_k(x, t - |x-X|/v_0)dX, \tag{42}$$

where the connectivity kernel $c_p(x,X)$ models the density of synapses of neurons whose soma is located at target location $x$, with afferent neurons having their somas at location $X$. The integral covers the spatial domain where units' somas can be found, and may thus be 1-, 2- or 3-dimensional depending on the model. The kernel $c_k$ reflects the spatial spread of axonal arborizations and as such can be approximated based on anatomical studies.

The elements finalizing the description of the state of the system are the relations between each population's membrane potential and rate, modeled by

$$\lambda_p = a_p(V_p)\,, \tag{43}$$

where $a_p$ is a typically sigmoidal activation function (these are the only non-linearities considered in our neural mass equations), leading to the overall dynamical system represented in Figure 5A.

[6] By "normalized", we mean that $s_k$ is a numerical quantity independent of the target population, possible target-specific differences in the PSCs being taken into account in the connectivity parameters $\nu_{p\leftarrow k}$, without loss of generality.

### From synaptic currents to LFPs

The local field potential is the lower frequency (<150Hz) part of the electrical potential recorded in the extracellular space, generated by the transmembrane currents [30]. Considering that active currents mostly reflect spiking activity, whose dynamics lies mostly above the typical LFP frequency range, we approximate the LFP as resulting from the linear superposition of passive membrane currents triggered by post-synaptic input currents [76], leading to the equation

$$L(y,t) = \sum_p \int f_{p,e}(y,x)\eta(x,t)dx + \sum_{p,k} \int f_{p,k}(y,x)s_k(x,t)dx\,,$$

where $f_{p,k}(.,x)$ represents the electrical field spatial distribution generated by trans-membrane currents of the $p$ cells with soma located at $x$, resulting from exciting them with post-synaptic unit currents of population $k$. Note due to charge neutrality of the cells, trans-membrane currents across the membrane of individual cells sum to zero, such that input currents at the levels of synapses are compensated by opposite trans-membrane currents away from them, typically leading to dipolar current distributions. Along the same lines, $f_{p,e}$ is the distribution associated with post-synaptic current resulting from exogenous inputs to $p$ cells. Differences between these fields according to the afferent populations are due to the respective distribution of their synaptic button over the efferent cell, preferentially targeting either the peri-somatic or distant dendritic sites, as illustrated in Figure 5C. These field distributions are assumed dominated by currents originating from pyramidal cells, due to their individual and collective geometric arrangement [82, 68, 76], such that we can simplify the above equation to obtain

$$L(y,t) = \int f_{E,e}(y,x)\eta(x,t)dx + \sum_k \int f_{E,k}(y,x)s_k(x,t)dx\,. \tag{44}$$

### Spike-LFP relation

Analysis of the frequency response of neural network models is a useful approach to understand their characteristics and underlying mechanisms [62, 107]. In our case, this can be performed analytically by linearizing Equation 43 around an operating point, the neural field model becomes a linear time-invariant system controlled by the exogenous input $\eta(t)$. We can thus compute transfer functions for each variable of the system that will determine their response to a sinusoidal exogenous input at frequency $f$, based on the computation of temporal Fourier transforms of the signals. For a given signal, $s(t)$ the temporal Fourier transform at frequency $f$ is given by

$$S(f) = \mathcal{F}_t[s](f) = \int_{\mathbb{R}} s(t)e^{-\boldsymbol{i}2\pi f t}dt\,.$$

By applying the Fourier transforms on the left- and right-hand-side of dynamical equations, we can derive transfer functions linking the responses of each network variable to the exogenous input at a given frequency. We provide in Table 3 a list of time-domain variables and their corresponding notations for their time-domain Fourier transform, used in analytical developments.

Specifically, for a general exogenous input signal with time-domain Fourier transform $\mathcal{E}(X,f)$, where $f$ denotes the temporal frequency, we obtain the input-output relation for population rates and LFP activity $L$

$$\lambda_p(x,f) = \int H_{\lambda_p}(x,X,f)\mathcal{E}(X,f)dX \quad \text{and} \quad L(y,f) = \int H_L(y,X,f)\mathcal{E}(X,f)dX\,.$$

A major simplification of this expression occurs when the exogenous input is separable in time and space,

$$\eta(X,t) = n(X)e(t))\,, \tag{45}$$

**Table 3.** List of neural field and mass model variables ($k$ indicates neuron population (afferent in case of synapse property))

| Description | Symbol | Temporal Fourier transform | Equation |
|---|---|---|---|
| Exogenous input (spatio-temporal) | $\eta(x,t)$ | $\mathcal{E}(x,f)$ | 41 |
| Exogenous input (temporal) | $e(t)$ | $\mathcal{E}(f)$ | 45 |
| Spike rate | $\lambda_k(x,t)$ | $\Lambda_k(x,f)$ | 46 |
| Membrane potential | $v_k(x,t)$ | $V_k(x,f)$ | 46 |
| Synaptic activity | $s_k(x,t)$ | $S_k(x,f)$ | 41, 42,44 |
| Activation function | $a_k(x,t)$ | $A_k(x,f)$ | 43 |

**Table 4.** List of neural mass model parameters

| Parameter name | Symbol | Value (*Mass2D*) | Value (*MassAlpha*) |
|---|---|---|---|
| $E$ membrane time constant | $\tau$ | $10ms$ | $10ms$ |
| $I$ membrane time constant | $\delta$ | $5ms$ | $10ms$ |
| $E \leftarrow I$ synaptic strength | $\nu_{E\leftarrow I}$ | 0.01 | 0.3 |
| $I \leftarrow E$ synaptic strength | $\nu_{I\leftarrow E}$ | 0.5 | 0.3 |
| Alpha synapse time constant | $\sigma$ | n.a. | $4ms$ |
| $I \leftarrow I$ synaptic strength | $\nu_{I\leftarrow I}$ | n.a. (accounted for in $\delta$) | 20.0 |

leading to $\mathcal{E}(X,f) = n(X)E(f)$ after temporal Fourier transform. This simplifying assumption models a number of typical inputs to the structure, including sinusoidal standing waves ($\eta(x,t) = n(x)e^{i2\pi ft}$) and traveling plane waves ($\eta(x,t) = e^{i(2\pi ft - kx)}$). This results in a simple expression for the covariance estimated across experimental trials between *rate* and LFP at two possibly different spatial points $(x, y)$

$$\langle L(y,f)\lambda_p(x,f)\rangle = \left(\int H_L(y,X,f)n(X)dX\right)\left(\int H_{\lambda_p}(x,X,f)n(X)dX\right)|e(f)|^2 .$$

in which the input intervenes only as a multiplicative positive constant, and which is separable in both space variables $x$ and $y$. As a consequence, the rank one approximation of the covariance between spiking units and LFP channels activity estimated by GPLA is informative about the microcircuit properties, as we explained in the Appendix describing the analysis of the neural mass and neural field models.

### Analysis and simulation of two population neural mass models

#### General description

The generic dynamic model of Equation 41 is exploited to describe network activity at a single location (i.e. we neglect the spatial extent of the considered structure) containing two cell types: pyramidal (E) and inhibitory (I), leading to the linear equations:

$$V_E + \tau_E \frac{dV_E}{dt} = \nu_{E\leftarrow E}s_E - \nu_{E\leftarrow I}s_I + \eta \quad (46)$$

$$V_I + \tau_I \frac{dV_I}{dt} = \nu_{I\leftarrow E}s_E - \nu_{I\leftarrow I}s_I + \alpha\eta \quad (47)$$

where $\boldsymbol{\nu}$ is a matrix gathering the non-negative synaptic strengths between populations, $\eta$ the exogenous input to the network, with $\alpha \geq 0$ controlling the ratio between feed-forward excitation and inhibition. The term $\nu_{kj}s_j$ is the population averaged post-synaptic potential from population $j$ to population $k$. In order to study quantitatively the effect of connectivity changes in the microcircuit, in this expression of the post-synaptic current, we isolate the synaptic strength coefficient $\nu_{k\leftarrow j}$, from a perisynaptic activity, that summarizes the dynamical processes occurring pre- and post-synaptically (synaptic delay, time constant induced by the post-synaptic channel conductance, ...). In the simplest case, we assume peri-synaptic activity $s_j$ can be approximated by the spike rate of population $j$, $\lambda_j$ (up to a multiplicative constant that is absorbed by $\nu_{kj}$). Alternatively, we model synaptic dynamics by a linear differential equation controlled by this rate (see model *MassAlpha* below).

The neural mass models will be analyzed with linear response theory, such that the $a_k$'s of Equation 43 will be linearized around an equilibrium point of the dynamical system (that can be computed for vanishing input $\eta = 0$), and the resulting multiplicative constants will be themselves absorbed in the connectivity matrix $\boldsymbol{\nu}$, leading to replacing Equation 43 by

$$\lambda_k = V_k\,. \tag{48}$$

Next we describe the two linearized neural mass models exploited to interpret GPLA results of hippocampal simulations (see Figure 6). Parameter values for both models are reported in Table 4

### *Mass2D*: E-I interactions without synaptic dynamics

Starting from Equation 46 and using Equation 48, the linearized system can be trivially reduced to the two-dimensional dynamical system (up to rescaling of the connectivity matrix coefficients)

$$\lambda_E + \tau\frac{d\lambda_E}{dt} = -\nu_{E\leftarrow I}\lambda_I + \eta\,, \tag{49}$$

$$\lambda_I + \delta\frac{d\lambda_I}{dt} = \nu_{I\leftarrow E}\lambda_E + \alpha\eta\,. \tag{50}$$

where $\tau$ and $\delta$ are time constants derived from membrane time constant $\tau_k$ and recurrent synaptic connection $\nu_{kk}$. Linear response analysis then relies on the Laplace transform (with Laplace variable $p$) of these equations

$$\Lambda_E(p) + \tau p\Lambda_E = -\nu_{E\leftarrow I}\Lambda_I + N(p)\,, \tag{51}$$

$$\Lambda_I(p) + \delta p\Lambda_I = \nu_{I\leftarrow E}\Lambda_E + \alpha N(p)\,. \tag{52}$$

For the case of *no feedforward inhibition* ($\alpha = 0$), this leads to the ratio of excitatory to inhibitory activity in the Laplace domain

$$\frac{\Lambda_E}{\Lambda_I}(p) = \frac{\delta p + 1}{\nu_{I\leftarrow E}} \tag{53}$$

resulting in excitatory activity being in advance of $\tan^{-1} 2\pi f\tau$ with respect to inhibitory activity at frequency $f$.

For the case of strong feedforward inhibition ($\alpha = 1$), this leads to

$$\frac{\Lambda_E}{\Lambda_I}(p) = \frac{\delta p + 1 - \nu_{E\leftarrow I}}{\tau p + 1 + \nu_{I\leftarrow E}} \tag{54}$$

such that the phase shift between the population is of constant sign across frequencies, but may be positive or negative depending on the exact parameters' values governing the E-I dynamics. Plots summarizing these situations are provided in Figure 6D.

### *MassAlpha*: E-I interactions with alpha type synaptic impulse response

Together with Equation 46, we include in addition a non-trivial synaptic dynamics in the form of the differential equation (assuming same dynamic for both AMPA and GABA synapses),

$$\sigma^2\frac{d^2 s_k}{dt^2} + 2\sigma\frac{ds_k}{dt} + s_k = \lambda_k\,, \tag{55}$$

This corresponds to the classical *alpha synapse* used in computational models (e.g. implemented in the *NEURON* software [18]), modeling the response to a single spike with the alpha function

$$s(t) = \frac{1}{\sigma^2} t e^{-t/\sigma} \,. \tag{56}$$

By combining these equations with linear activations (Equation 48), the dynamics of the circuit is summarized by a 6-dimensional state-space model that can be studied analytically with linear response theory.

### Analysis and simulations of neural field models

When taking into account the spatial extension of the network, neural mass models can be extended to neural field models, where the variables described above possibly depend on space. Consider one or two spatial dimensions tangential to the layers of the network (assuming a layered organization like the hippocampus or cortex), the key phenomenon that should be additionally modeled is then the coupling between activity in different locations of the network entailed by horizontal connections. In line with the literature and to simplify the analysis, we will consider only the excitatory connections are spatially extended. With respect to the above generic neural field model Equations 41-43, equations pertaining to synaptic activity and rate need only to be specified. We use the model introduced by Jirsa and Haken [55, Equation 15] with a spatial diffusion term with characteristic distance $r_0$ and axonal propagation speed $v_0 = r_0\gamma$, which takes the form of a damped wave equation:

$$\frac{1}{\gamma^2}\frac{\partial^2 s_E}{\partial t^2} + \frac{2}{\gamma}\frac{\partial s_E}{d\partial t} + s_E - r_0^2 \Delta s_E = \lambda_E + \frac{1}{\gamma}\frac{\partial \lambda_E}{\partial t} \,, \tag{57}$$

where $\Delta$ is the Laplacian operator, while $s_I = V_I$ to encode purely local inhibition, eliminating redundant multiplicative factors.

To specify Equation 43, we use sigmoid activation functions for both AMPA and GABA synapses,

$$\lambda_E = \frac{Q_E}{1+\exp(-\chi_E.(V_E - V_{th,E}))} \,, \tag{58}$$

$$\lambda_I = \frac{Q_I}{1+\exp(-\chi_I.(V_I - V_{th,I}))} \,, \tag{59}$$

whose parameters (maximum rate $Q_k$, spiking threshold $V_{th,k}$ and excitability $\chi_k$ for population $k$) are adjusted in order to obtain different types of dynamics, either evolving around a stable equilibrium point (model *FieldStable*) or with a clear oscillatory activity (model *FieldOsc*).

#### Spatio-temporal phase analysis in 1D

Before simulating 2D neural field models with an explicit method, we investigate analytically properties of a simplified 1D model. In this case, the partial differential Equation 57 corresponds to an exponentially decaying connectivity, with axonal propagation speed $v_0 = r_0\gamma$, such that the resulting post-synaptic current takes the integral form (see Jirsa and Haken [55, Equation (14)])

$$s_E(x,t) = \frac{1}{2r_0}\int \exp(-|x-X|/r_0)\lambda_E(X, t-|x-X|/v_0)dX \,. \tag{60}$$

If we take the neural field equation in the context of horizontal connection along non-myelinated axons ($v_0 \sim 1m/s$, $r_0 < 1mm$), the typical value of $\gamma$ is beyond 1000 such that if we focus on frequencies below $200Hz$, we may neglect for the temporal derivatives of the partial differential equation. This leads to the following approximation for the dynamics of excitatory post-synaptic current.

$$s_E - r_0^2 \frac{\partial^2 s_E}{dx^2} = \lambda_E \,. \tag{61}$$

In an unbounded 1D medium, assuming that activities vanish at large distances, we can use the spatial Fourier transform $\widehat{f}(t,z) = \mathcal{F}_x\left[f(t,x)\right](z) = \int_{\mathbb{R}} f(t,x)e^{-2\boldsymbol{i}\pi z x}dx$ to derive the expression of $s_E$ as a function of $\lambda_E$

$$\widehat{s_E}(t,z) = \frac{1}{1+(2\pi z)^2 r_0^2}\widehat{\lambda_E}(t,z)\,. \tag{62}$$

In order to get back to the original spatial position domain, we use a general Fourier transform relation for an arbitrary complex parameter $a$ such that $\mathrm{Re}[a] \geq 0$:

$$\mathcal{F}_x\left[\frac{1}{2a}e^{-|x|a/r_0}\right](z) = \frac{r_0}{(2\pi z r_0)^2 + a^2}\,, \tag{63}$$

This formula can be inverted, considering an arbitrary complex number $b$, and defining $\sqrt{b}$ to be the unique complex number such that $\sqrt{b}^2 = b$ and $\mathrm{Re}\sqrt{b} \geq 0$, we get

$$\mathcal{F}_z^{-1}\left[\frac{r_0}{(2\pi z)^2 r_0^2 + b}\right](x) = \frac{1}{2\sqrt{b}}e^{-|x|\sqrt{b}/r_0}\,, \tag{64}$$

For the particular case $b=1$, we get $\sqrt{b}=1$, which, in the spatial position domain, leads to

$$s_E(t,x) = \int_{\mathbb{R}} \lambda_E(t,y)h_{r_0}(x-y)dy = h_{r_0} * \lambda_E(t,x)\,, \tag{65}$$

where $*$ denotes spatial convolution and $h_{r_0}(x) = \frac{1}{2r_0}e^{-|x|/r_0}$. This reflects that horizontal connectivity generates EPSCs corresponding to a spatial smoothing of the excitation rate spatial distribution.

After linearizing around the operating point of the network (absorbing again the resulting multiplicative constant in the connectivity matrix), we obtain the equation of the dynamics by modifying Equation 41 (assuming neither long range nor feedforward inhibition)

$$\lambda_E(t,x) + \tau\frac{d\lambda_E}{dt} = \nu_{E\leftarrow E}s_E(t,x) - \nu_{E\leftarrow I}\lambda_I(t,x) + \eta(t,x)\,, \tag{66}$$

$$\lambda_I(t,x) + \delta\frac{d\lambda_I}{dt} = \nu_{I\leftarrow E}s_E(t,x)\,, \tag{67}$$

where the synaptic strength values incorporate multiplicative constants resulting from the linearization of Equations 58-59. By computing the temporal (with frequency variable $f$) and spatial Fourier transform of each equation, we get (using $p = \boldsymbol{i}2\pi f$ )

$$\widehat{\Lambda_E} + \tau p\widehat{\Lambda_E} = \nu_{E\leftarrow E}\widehat{S_E} - \nu_{E\leftarrow I}\widehat{\Lambda_I} + \widehat{H}\,, \tag{68}$$

$$\Lambda_I(t,x) + \delta p\widehat{\Lambda_I} = \nu_{I\leftarrow E}\widehat{S_E}\,. \tag{69}$$

Eliminating $\widehat{\Lambda_I}$ we get

$$(1+\tau p)\widehat{\Lambda_E} = \left(\nu_{E\leftarrow E} - \nu_{E\leftarrow I}\frac{\nu_{I\leftarrow E}}{1+\delta p}\right)\widehat{S_E} + \widehat{H}\,. \tag{70}$$

Combined with the spatially Fourier transformed horizontal connectivity Equation 61 (using $k = \boldsymbol{i}2\pi z$)

$$(1 - r_0^2 k^2)\widehat{S}_E = \widehat{\Lambda_E}\,, \tag{71}$$

this leads to

$$\widehat{S_E} = \frac{1}{1+\tau p} \frac{\widehat{E}}{-r_0^2 k^2 + 1 + \frac{1}{1+\tau p}\left(\frac{\nu_f}{1+\delta p} - \nu_{E\leftarrow E}\right)}\,, \tag{72}$$

where we define the *feedback inhibition gain* $\nu_f = \nu_{E\leftarrow I}\nu_{I\leftarrow E}$.

Introducing our time-space separability assumption on the exogenous input

$$\eta(x,\, t) = n(x)\epsilon(t) \tag{73}$$

leads to

$$\widehat{S_E}(z,\, f) = \frac{1}{1+\boldsymbol{i}2\pi\tau f} \frac{E(f)\widehat{n}(z)}{-r_0^2(2\pi z)^2 + 1 + \frac{1}{1+\boldsymbol{i}2\pi\tau f}\left(\frac{\nu_f}{1+\boldsymbol{i}2\pi\delta f} - \nu_{E\leftarrow E}\right)}\,, \tag{74}$$

By defining

$$b = 1 + \frac{1}{1+\boldsymbol{i}2\pi\tau f}\left(\frac{\nu_f}{1+\boldsymbol{i}2\pi\delta f} - \nu_{E\leftarrow E}\right)\,, \tag{75}$$

and using the inverse spatial Fourier transform of Equation 64, we get

$$S_E(x,\, f) = \frac{1}{2r_0\sqrt{b}} \frac{E(f)}{1+\boldsymbol{i}2\pi\tau f} n(x) * e^{-|x|\sqrt{b}/r_0}\,, \tag{76}$$

Assuming the exogenous input does not impose a spatial phase gradient to the structure (i.e. $n(x)$ is positive real for all locations up to a multiplicative constant), the phase gradient at a given frequency will be controlled by the imaginary part of $\sqrt{b}$. Specifically, to investigate qualitatively the phase gradient around a peak of activity of the exogenous input, we assume that $n(x)$ is a dirac at $x = 0$. Then the spatial variation of the phase around $x = 0$ take the form

$$\phi(x) = -\frac{|x|}{r_0}\mathrm{Re}\left[\sqrt{b}\right]\,. \tag{77}$$

This dirac approximation, does not match well our simulations (using a Gaussian shape spatial input distribution). However, computing spiking activity form $S_E$ based on Equation 61, to obtain the spatial distribution of the spike vector, will have a deblurring effect compensating the convolution by $n(x)$ in Equation 76, making in closer to a Dirac. As a consequence, we will interpret the data based on the following approximation

$$\lambda_E(x,\, f) \approx C\frac{1}{2r_0\sqrt{b}} \frac{E(f)}{1+\boldsymbol{i}2\pi\tau f} e^{-|x|\sqrt{b}/r_0}\,, \tag{78}$$

up to a multiplicative constant $C$.

In order to investigate the qualitative effect of the microcircuit connectivity on this spatial gradient, we assume $\tau = \delta$ and use a low (temporal) frequency assumption of the form $f << 1/\tau$, such that we can exploit a first order expansion for the fractions containing the term $\tau p << 1$. This leads to the approximation

$$b \approx 1 + (1 - \boldsymbol{i}2\pi\tau f)\left(\nu_f(1 - \boldsymbol{i}2\pi\tau f) - \nu_{E\leftarrow E}\right) \approx 1 + \nu_f - \nu_{E\leftarrow E} - \boldsymbol{i}2\pi\tau f(2\nu_f - \nu_{E\leftarrow E})\,. \tag{79}$$

Simple geometric considerations show that the sign of the imaginary part of $b$ is the same as the sign of its square root, such that under our simplifying assumption, the sign of the gradient taken algebraically from center ($x = 0$) to surround ($|x| > 0$) is the sign of

$$2\nu_f - \nu_{E\leftarrow E}\,, \tag{80}$$

showing that strong feedback inhibition will tend to put the populations surrounding $x = 0$ in advance with respect to this center point, while weak feedback inhibition (with respect to feedback excitation), will to generate a phase lag of the surround with respect to the center.

### Neural field simulation in 2D

While the above analysis is much easier to perform in 1D, in most structures (and in particular cortex), the domain spanned by horizontal connectivity is better approximated by a 2D domain, which can also be sampled by modern electrode arrays. We thus simulate the dynamics of such 2D system to get insight into the characteristics revealed by GPLA analysis in this context.

The equations of the continuous field dynamics are non-linear, comprising Equations 57, 58 and 59 as well as the following membrane dynamics for each population

$$V_E(t,x,y) + \tau \frac{dV_E}{dt} = \tilde{\nu}_{E\leftarrow E} s_E(t,x,y) - \tilde{\nu}_{E\leftarrow I}\lambda_I(t,x,y) + \eta(t,x,y)\,, \tag{81}$$

$$V_I(t,x,y) + \delta \frac{dV_I}{dt} = \tilde{\nu}_{I\leftarrow E} s_E(t,x,y)\,, \tag{82}$$

where we use the modified notation $\tilde{\nu}_{Q\leftarrow P}$ for the synaptic strengths, such that we can keep the notation $\nu_{Q\leftarrow P}$ to describe the synaptic strengths of the system linearized around its operating point (values at this point are denoted with superscript $^{op}$ in the following). The linearizations of Equations 58 and 59 then entails

$$\nu_{E\leftarrow E} = \tilde{\nu}_{E\leftarrow E}\lambda_E^{op}(1-\lambda_E^{op}/Q_E)\chi_E\,, \tag{83}$$

$$\nu_{I\leftarrow E} = \tilde{\nu}_{I\leftarrow E}\lambda_E^{op}(1-\lambda_E^{op}/Q_E)\chi_E\,, \tag{84}$$

$$\nu_{E\leftarrow I} = \tilde{\nu}_{E\leftarrow I}\lambda_I^{op}(1-\lambda_I^{op}/Q_I)\chi_I\,. \tag{85}$$

where expressions exploit the fact that the derivative of the used sigmoid function $\sigma(x) = \frac{1}{1+e^{-x}}$ is $\frac{d\sigma}{dx}(x) = \sigma(x)\sigma(1-\sigma(x))$. These synaptic strengths of the linearized system are used in Supplementary Figure 6 compare the phase-modulus relation of the spike vector with theoretical predictions based on Equation 9.

We use simplified notations for the 2D (in space) time-varying scalar fields $V(t,x,y) = s_E((x,y),t)$ and $I(t,x,y) = \lambda_E((x,y),t)$. Let $\Delta x$ and $\Delta t$ be the spatial and temporal grid spacing, and $V_{j,l}^n = V(n\Delta t, j\Delta x, l\Delta x)$ the discretized field. We use a Forward Time Centered Space (FTCS) finite difference scheme to simulate the above neural field model [38]. FTCS relies on making the approximations

$$\frac{\partial V}{\partial t}(t,x,y) \approx \frac{1}{\Delta t}\left(V_{j,l}^{n+1} - V_{j,l}^{n}\right)\,, \quad \frac{\partial^2 V}{\partial t^2}(t,x,y) \approx \frac{1}{(\Delta t)^2}\left(V_{j,l}^{n+1} + V_{j,l}^{n-1} - 2V_{j,l}^{n}\right)\,, \tag{86}$$

$$\text{and} \quad \frac{\partial^2 V}{\partial x^2}(t,x,y) \approx \frac{1}{(\Delta x)^2}\left(V_{j+1,l}^{n} + V_{j-1,l}^{n} - 2V_{j,l}^{n}\right)\,. \tag{87}$$

Applying these approximations to Equation 57, leads to an explicit scheme for the field values at time $(n+1)\Delta t$ based on all values at time $n\Delta t$ and $(n-1)\Delta t$.

$$V_{j,l}^{n+1} = \frac{1}{G+1}\left((G-1)V_{j,l}^{n-1} + 2V_{j,l}^{n} + G^2\left(R^2\left(K_\ell *_d \boldsymbol{V}^n\right)_{j,l} + I_{k,l}^{n} + \frac{1}{G}\left(I_{k,l}^{n} - I_{k,l}^{n-1}\right) - V_{j,l}^{n}\right)\right)\,, \tag{88}$$

where $G = \gamma\Delta t$, $R = r_0/\Delta x$ and $K_\ell *_d$ denotes the discrete 2D spatial convolution with the discrete Laplace operator

$$K_\ell = \begin{bmatrix} 0 & 1 & 0 \\ 1 & -4 & 1 \\ 0 & 1 & 0 \end{bmatrix}\,. \tag{89}$$

The parameters chosen for both models presented in the main text are reported in Table 5.

**Table 5.** List of 2D neural field model parameters

| Parameter name | Symbol | Value *(for each level of recurrent inhibition)* | | | |
|---|---|---|---|---|---|
| | | *weak* | *lower med.* | *upper med.* | *strong* |
| $E$ membrane time constant | $\tau_E$ | $20ms$ | $20ms$ | $20ms$ | $20ms$ |
| $I$ membrane time constant | $\tau_I$ | $20ms$ | $20ms$ | $20ms$ | $20ms$ |
| E-E synaptic strength | $\tilde{\nu}_{E\leftarrow E}$ | 0.2 | 0.2 | 0.2 | 0.2 |
| I-I synaptic strength | $\tilde{\nu}_{I\leftarrow I}$ | 0 | 0 | 0 | 0 |
| $E \rightarrow I$ synaptic strength | $\tilde{\nu}_{E\leftarrow I}$ | 0.2 | 0.2 | 0.2 | 0.2 |
| $I \rightarrow E$ synaptic strength | $\tilde{\nu}_{I\leftarrow E}$ | 1 | 1 | 1 | 1 |
| $E$ excitability | $\chi_E$ | 1 | 1 | 1 | 1 |
| $I$ excitability | $\chi_I$ | 0.1 | .33 | 1 | 3.33 |
| $E$ sigmoid threshold | $V_{th,E}$ | 0 | 0 | 0 | 0 |
| $I$ sigmoid threshold | $V_{th,I}$ | 0 | 1 | 5 | 5 |
| $E$ maximum rate | $Q_E$ | $20Hz$ | $20Hz$ | $20Hz$ | $20Hz$ |
| $I$ maximum rate | $Q_I$ | $20Hz$ | $20ms$ | $20ms$ | $20Hz$ |

### Quantification and statistical analysis

#### Parameter estimation of von Mises distribution

The von Mises distribution (VM), which is also known as "circular normal" distribution is the counterpart of the Gaussian distribution for circular data [37, Chapter 3]. We used it for various purposes in this work (e. g. to model the spiking probability to synthesize phase-locked spike trains).

The VM distribution takes the form,

$$p(\phi|\varphi_0, \kappa) = \frac{1}{2\pi I_0(\kappa)} \exp\left(\kappa \cos(\phi - \varphi_0)\right), \tag{90}$$

where $I_0(\kappa)$ is the modified Bessel function of order zero (Equation 36). In Figure 8 and 7, we fit a VM distribution to the pooled phases of spike and LFP vectors coefficients. We use a maximum likelihood (ML) method for estimating the two parameters of the VM distribution, $\varphi_0$ and $\kappa$ [37]. The ML estimation of $\varphi_0$ is simply the sample mean direction, denoted by $\bar{R}$ (for spike-LFP data, is the locking phase). Maximum likelihood estimation of, $\widehat{\kappa}$, is the solution of following equation:

$$A_1(\widehat{\kappa}) = \bar{R}, \tag{91}$$

and $A_1$ is a ratio of two modified Bessel functions:

$$A_1(x) = \frac{I_1(x)}{I_0(x)}. \tag{92}$$

Approximate solutions are available for $\widehat{\kappa}$ [37, sec. 4.5.5]

$$\widehat{\kappa} = \begin{cases} 2\overline{R} + \overline{R}^3 + 5\overline{R}^5/6 & \overline{R} < 0.53 \\ -0.4 + 1.39\overline{R} + 0.43/(1-\overline{R}) & 0.53 \le \overline{R} < 0.85 \\ 1/\left(\overline{R}^3 - 4\overline{R}^2 + 3\vec{R}\right) & \overline{R} \ge 0.85 \end{cases} \tag{93}$$

where $\overline{R}$ is the resultant length of the phases.

#### Computing Signal-to-Noise Ratio

In Figure 3H-I, in order to compare GPLA-based and PLA-based estimation of pairwise couplings, we define signal-to-Noise Ratio (SNR) as the ratio of coupling strength (PLV) to estimation error (the difference between estimated PLV and the ground truth) and it was used to compare the quality of GPLA-based and univariate estimation.

For the PLA-based estimation, SNR is the ratio of PLV to its estimation error. To compute the SNR for the GPLA-based estimation, the following procesure has been used: 1- Compute PLVs for all pairs of spiking units and LFP signals (based on Equation 16). 2- Construct the coupling matrix with the obtained PLVs, as shown in Figure 3G. 3- Obtain the rank-one approximation of the coupling matrix via GPLA (which is based on SVD at core). 4- Compare the elements of the approximated coupling matrix to the ground truth PLV, and compute the SNR similar to PLA-based computation.

### Animal preparation and intracortical recordings

The methods for surgical preparation, anesthesia, and presentation of visual stimuli for the Utah array recordings have been described in previous studies (see [70, 69, 5, 102]).

#### Data collection

Neural signals were recorded with a NeuroPort Cortical Microelectrode Array (Blackrock Microsystems, Salt Lake City, Utah USA). An array was implanted in the inferior convexity of the prefrontal cortex (see [102] for more details). The arrays are 4mm $\times$ 4mm with a 10 by 10 electrode configuration. Neural signals recorded from 96 of the available 100 electrodes. Neural activity was recorded in 200 trials. Each trial consisted of a 10s period of movie presentation, followed by 10s of a blank screen (inter-trial).

#### LFP extraction

The raw signals were low-pass filtered using an 8th order Chebyshev Type 1 filter with a cut-off frequency of 200Hz and a pass-band ripple less than 0.05dB. Forward and backward filtering was used to minimize phase distortions caused by the filtering. Next, the filtered signal was decimated to a sampling frequency of approximately 500Hz.

#### Spike detection

For detecting multi-unit spikes, the raw signal was band-pass filtered using a minimum–order finite impulse response (FIR) filter [94] with pass-band cut-off frequencies of 600Hz to 5800Hz and stop-band cut-off frequencies of 400Hz and 6000Hz, with at least 65dB attenuation in the stop-bands and less than 0.002dB ripple within the pass-band. The amplitude threshold for spike detection was set to 5 standard deviations above the average of the filtered signal [93]. To spare computational costs, the standard deviation of the signal for each channel was estimated using a smaller, randomly chosen section of the filtered signal. Spike times with inter-spike intervals less than the refractory period of 0.5ms were eliminated.

1825 # Supplementary Figures

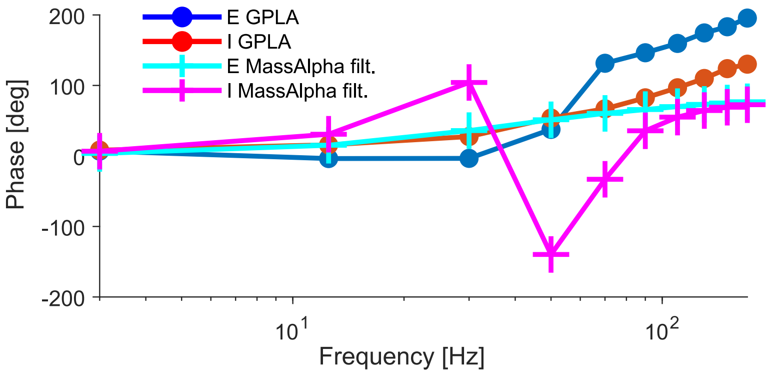


**Supplementary Figure 1. Use of EPSP as LFP proxy**.
Difference between phase of excitatory and inhibitory neurons/populations based on GPLA and the excitatory and inhibitory populations in the MassAlpha neural mass model. In this simulation EPSP has been used for the LFP proxy.

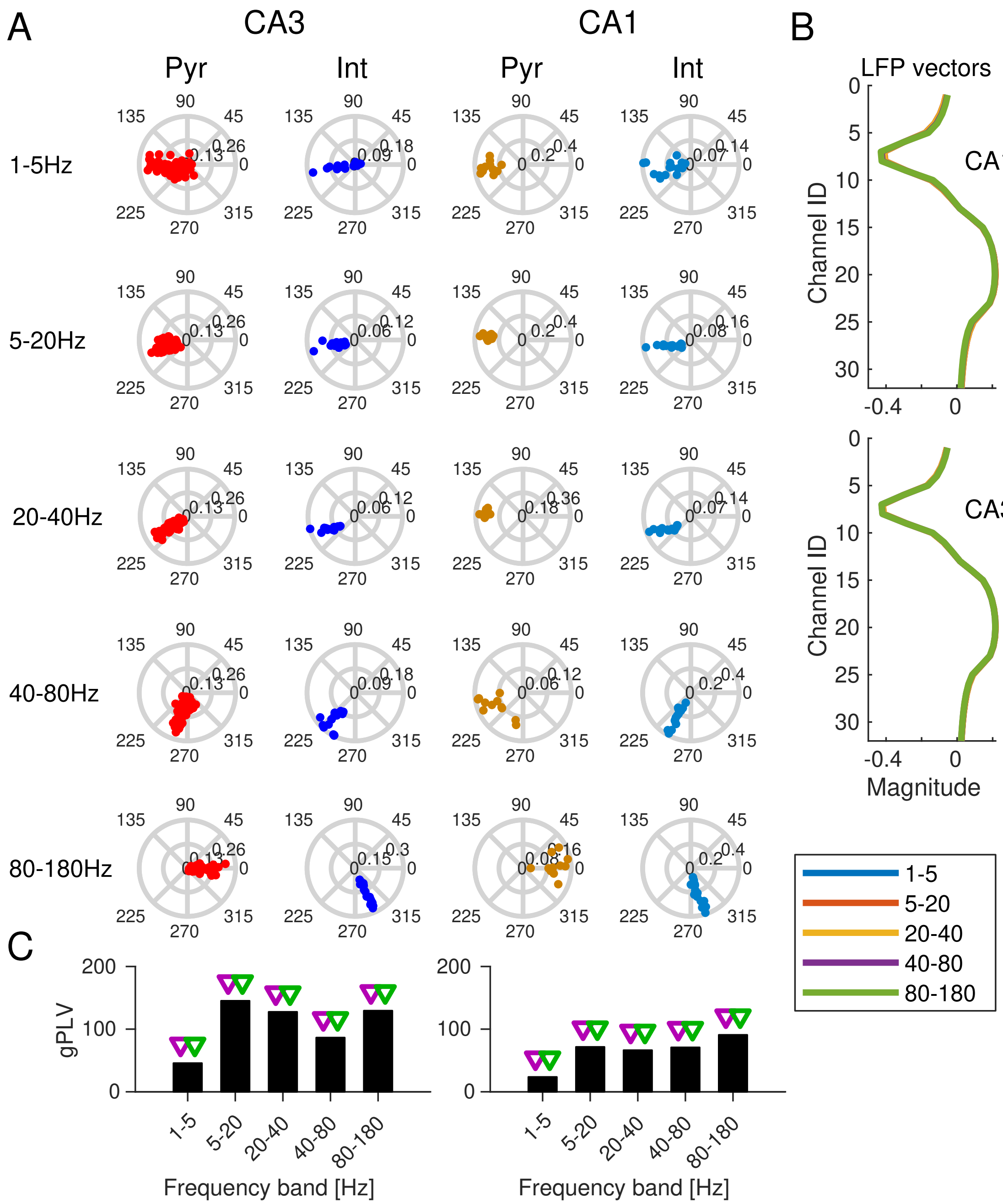


**Supplementary Figure 2. Joint GPLA of CA3 and CA1 activities.**
For this analysis, CA1 and CA3 data were separately injected to GPLA. **(A)** Spike vectors represented in polar plots similar to Figure 6E, but for all frequencies (indicated on the left). **(B)** LFP vectors, similar to Figure 6D, but for all frequencies (indicated in legend in the bottom). **(C)** gPLV for different frequency ranges of LFPs, similar to Figure 6C. Triangles indicated the significance assessed based on empirical (blue triangles, with significance threshold of 0.05) and theoretical (red triangles) tests. (left) for CA3 and (right) for CA1.

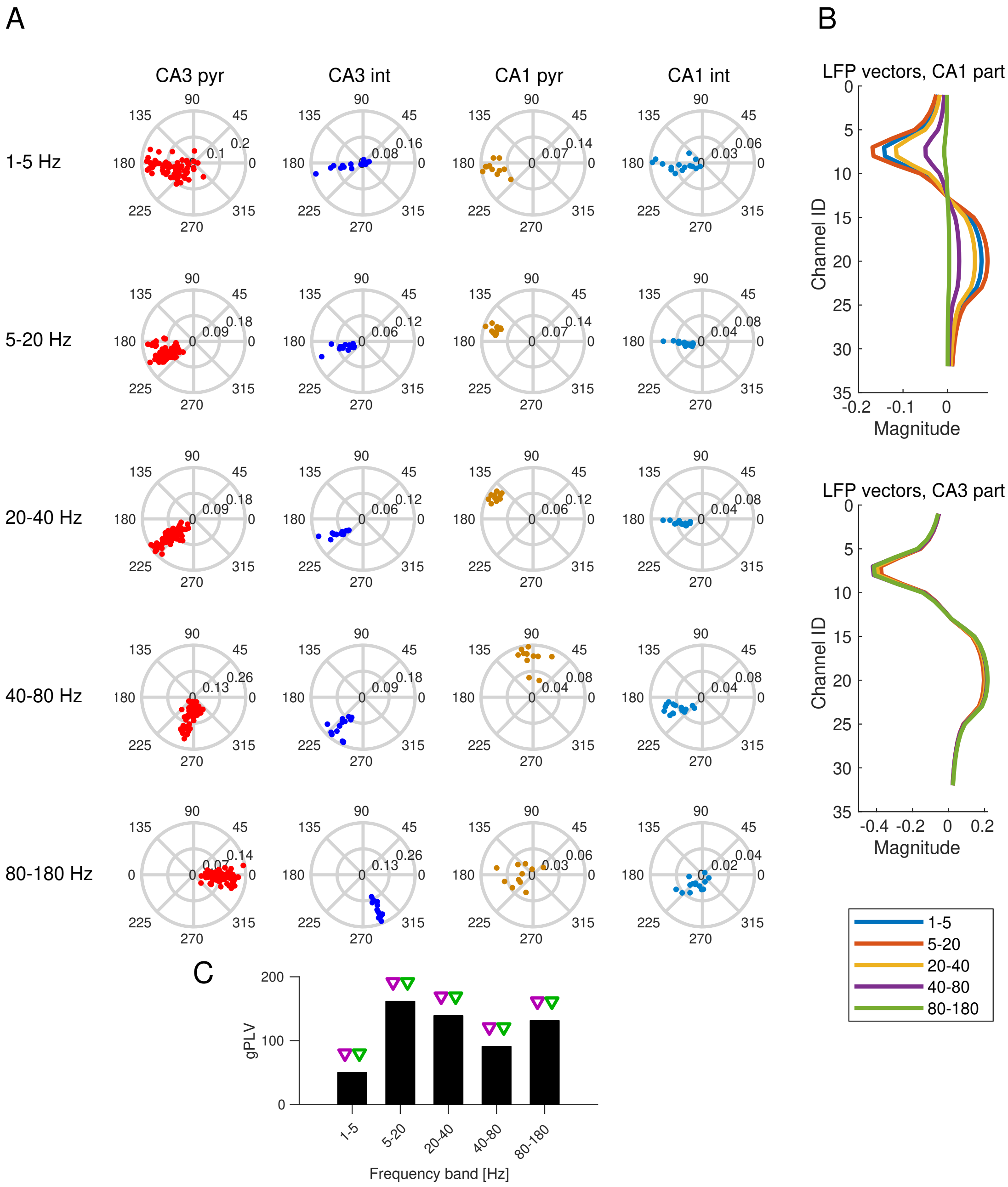


**Supplementary Figure 3. Joint CA1-CA3 analysis of hippocampal SWRs generated by a computational model**

For this analysis, CA1 and CA3 data were injected to GPLA together. **(A)** Spike vectors represented in polar plots similar to Figure 6E, but for all frequencies (indicated on the left). **(B)** LFP vectors, similar to Figure 6D, but for all frequencies (indicated in legend in the bottom). **(C)** gPLV for different frequency ranges of LFPs Figure 6C. Triangles indicated the significance assessed based on empirical (blue triangles, with significance threshold of 0.05) and theoretical (red triangles) tests.

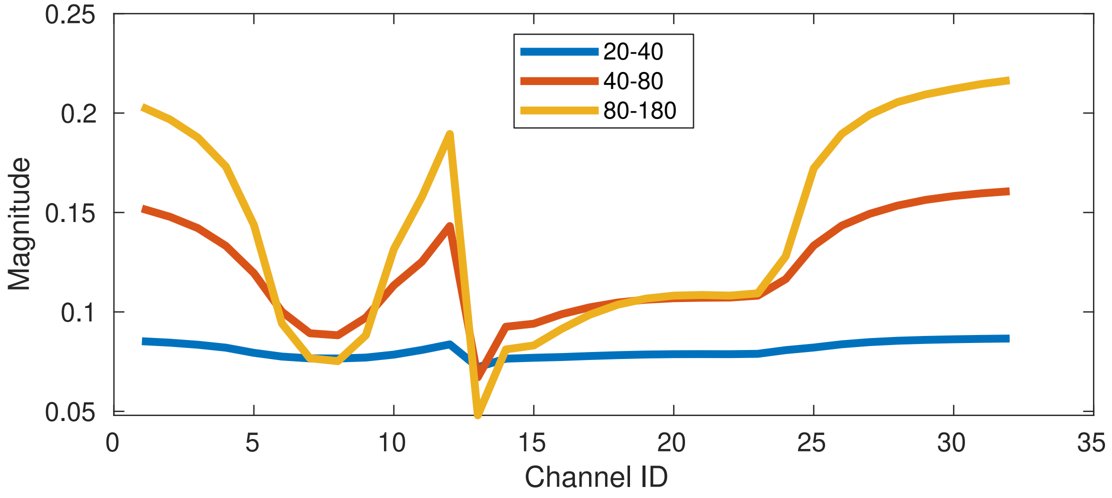


**Supplementary Figure 4. GPLA vs. PLA comparison for hippocampal SWR simulation.**
Similar to Figure 6D but based on uni-variate phase locking analysis (rather than multivariate GPLA). Each line depicts the phase locking value (PLV) for a fixed spiking units across all LFP channels. Colors indicate the frequency of filtered LFP.

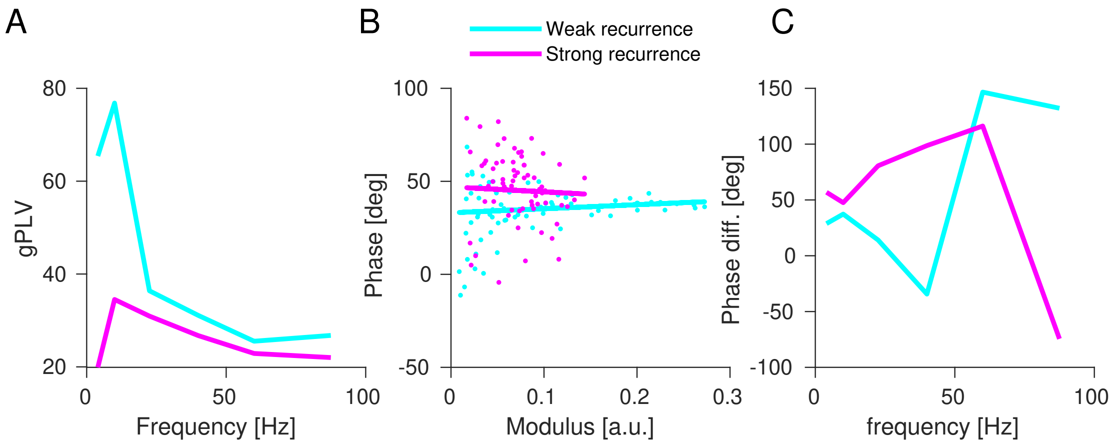


**Supplementary Figure 5. GPLA using IPSP as LFP proxy.**
To be compared with Figure 7C-E. **(A)** gPLV as a function of frequency for both models. **(B)** Phase of spike vector coefficients as a function of its modulus for the frequency band associated with maximum gPLV for each model (each dot one coefficient, and the continuous lines are plotted based on linear regression). **(C)** Shift between the averaged phase of spike vector and averaged phase of LFP vector, as a function of frequency.

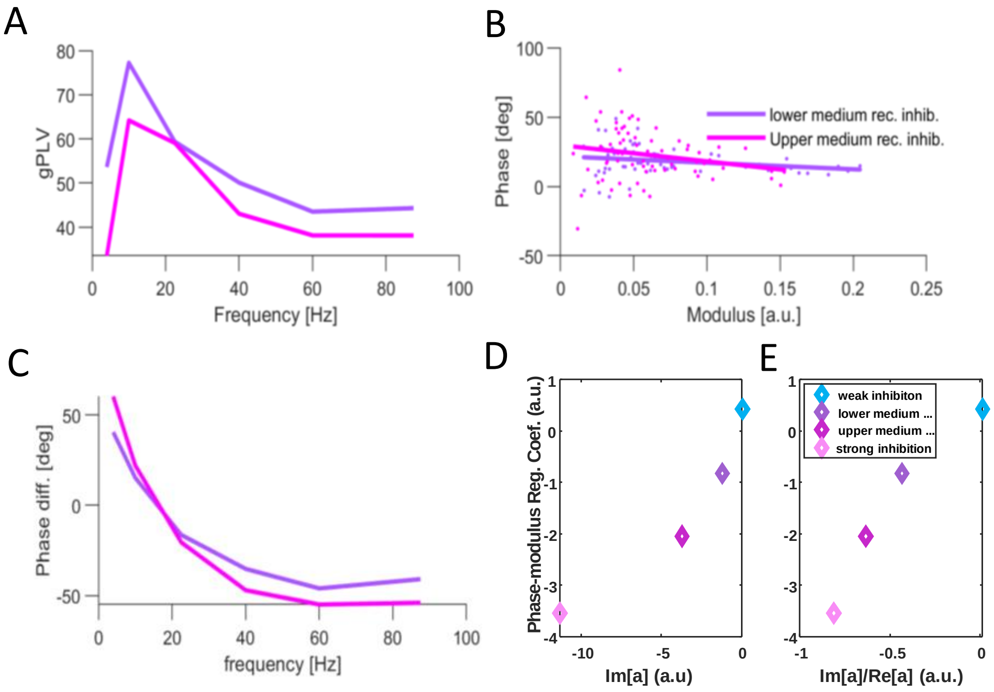


**Supplementary Figure 6. Phase-modulus relation dependency on level of inhibition.** Related to Figure 7D. **(A)** Same as Figure 7C. for simulations at intermediate levels of recurrent inhibition. **(B)** Same as Figure 7D. for simulations at intermediate levels of recurrent inhibition. **(C)** Same as Figure 7E. for simulations at intermediate levels of recurrent inhibition. **(D)** Magnitude of phase modulus regression coefficient (rescaled by $180/\pi$ to have it in radians) as a function of imaginary part of $a$ derived from Equation 9. **(E)** Same as (A) for $\text{Im}[a]/\text{Re}[a]$ instead of $\text{Im}[a]$.

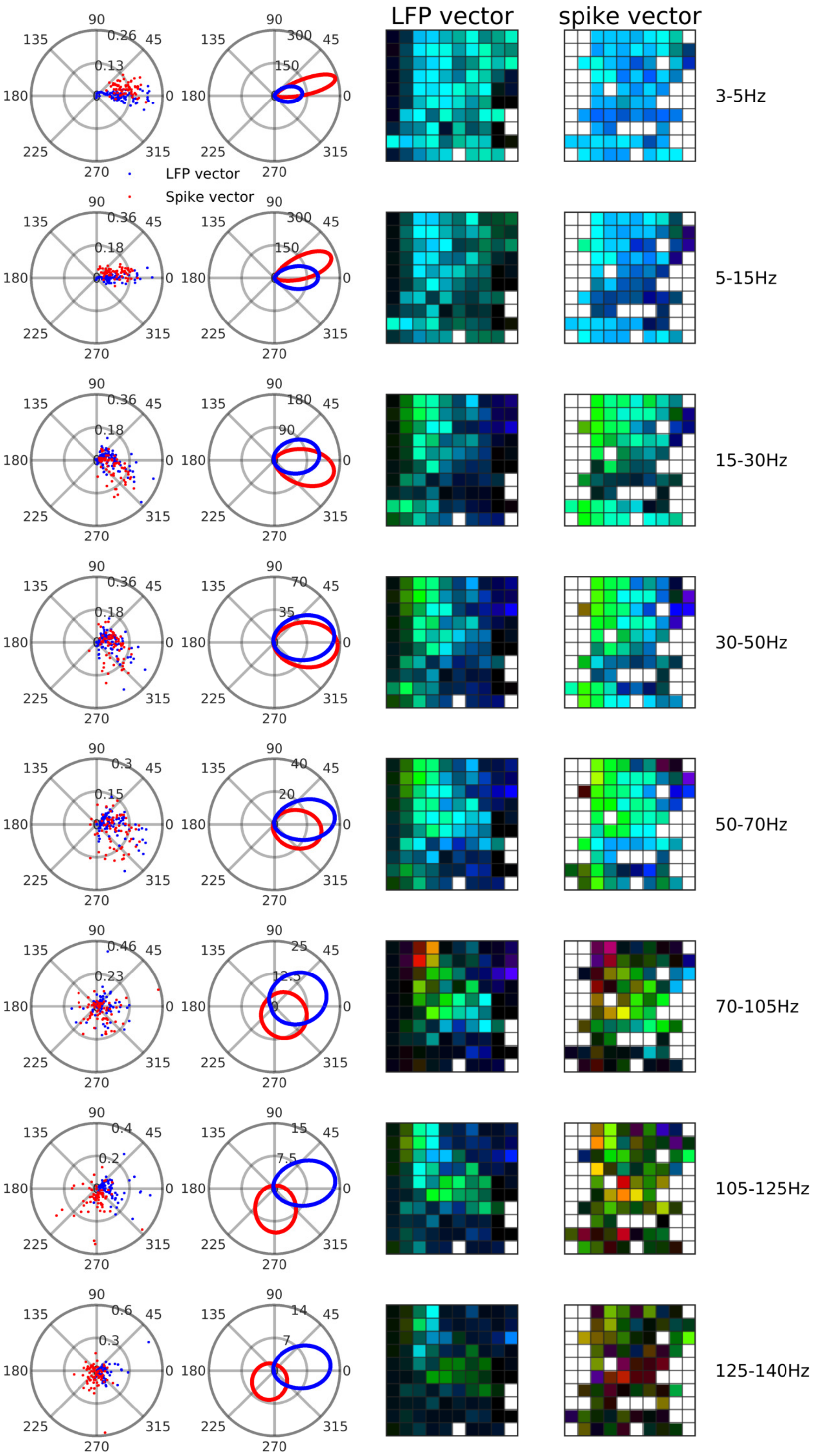


**Supplementary Figure 7. Analysis of PFC Utah array data**

LFP and spike vectors for frequencies indicated on the right. First column depict the LFP (blue dots) and spike (red dots) in the complex plane. Second column depict fitted von Mises distribution to phase of LFP and spike vectors. Third and forth column respectively represnting phase of LFP and spike vectors which remapped to real configuration of electrodes on Utah array (see Figure 8C).

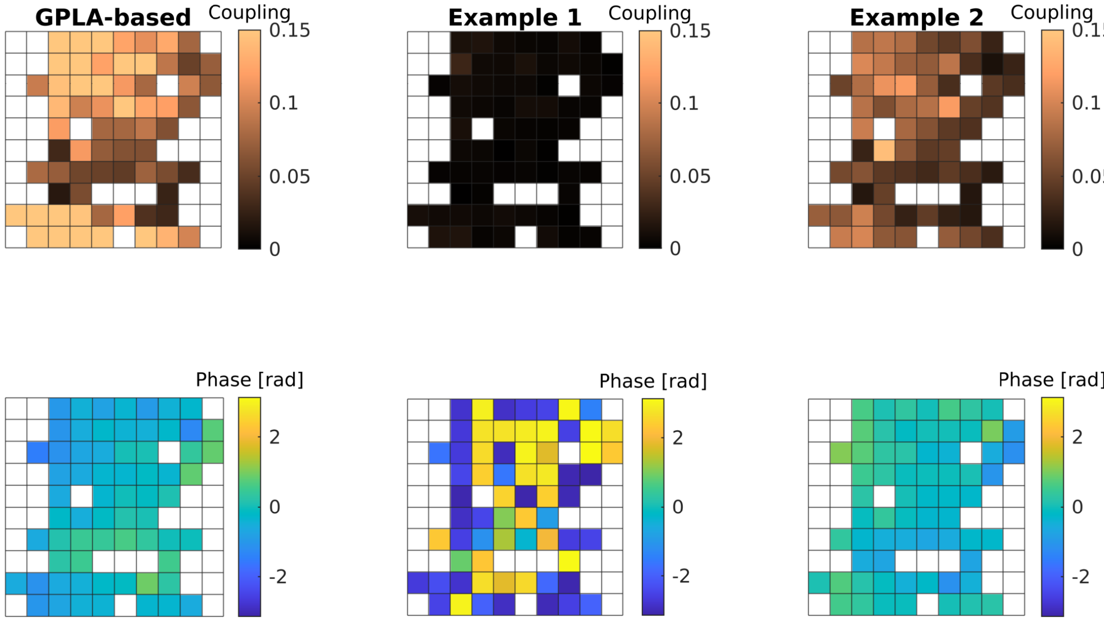


**Supplementary Figure 8. GPLA vs. PLA comparison for PFC Utah array data in revealing spatial pattern of coupling.**

Similar to Utah array maps in Figure 8I but based on uni-variate phase locking analysis (rather than multivariate GPLA). Panels in the first row depict the spatial distribution of phase locking value (PLV) or magnitude of the spike-field coupling on the array (see Figure 8C ). Panels in the second row depict the spatial distribution of locking phase on the array. White pixels in all panels indicate the recording channels with insufficient number of spikes (multiunit activity with a minimum of 5 Hz firing), as it was used in Figure 8I . The colorbars indicate the couple strength in the first row; and locking phase in the second row. First column, depicts the results based on multivariate GPLA, and second and third column depicts the results based on uni-variate phase locking analysis, but for two different choice of LFP reference channel. The result from 'Example 2' is close to what is captured based on GPLA, however result from 'Example 1' does not, due to lack of global coupling.

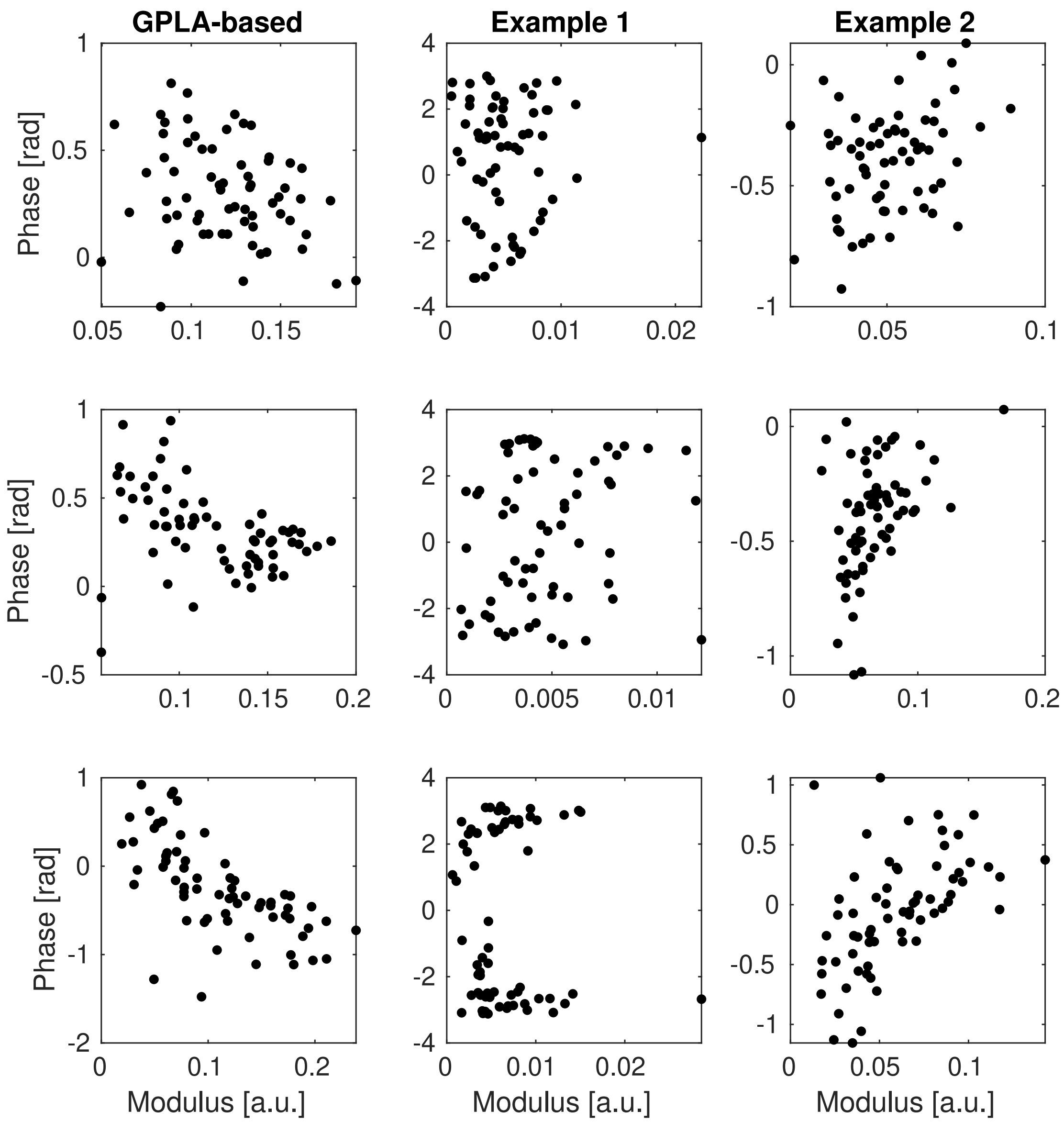


**Supplementary Figure 9. GPLA vs. PLA comparison for PFC Utah array data in characterizing strength of recurrent inhibition in PFC circuit.**
Similar to Figure 8H but based on uni-variate phase locking analysis (rather than multivariate GPLA). Each row correspond to analysis in different frequency (the same frequencies used in Figure 8H ), i. e. , 3-5 Hz, 5-15 Hz, and 15-30 Hz, respectively, first, second and third row. First column indicates the results based on GPLA (notebely pairwise coupling measure used here is exactly PLV), and imply the negative slope, similar to Figure 8H. The second and third columns demonstrate a similar analysis based on phase locking analysis, i. e. , locking phase plotted versus strength of coupling (PLV). Two example of LFP reference channels (the same references used in Figure supplement 8 ) Notably, none are compatible with our mean-field analysis (Figure 7) , thus not conclusive about the strength of recurrent inhibition.
